# Global plate model choice impacts reconstructions of the latitudinal biodiversity gradient

**DOI:** 10.1101/2025.01.09.632144

**Authors:** Lewis A. Jones, William Gearty, Lucas Buffan, Bethany J. Allen

## Abstract

The latitudinal gradient of declining species richness from the Equator towards the poles is one of the most pervasive macroecological patterns on Earth today. However, the ubiquity of this trend over geological timescales remains unclear. One reason for this uncertainty is that palaeobiologists need Global Plate Models (GPMs) to estimate the latitudinal position of organisms’ remains at time of deposition. However, as GPMs constitute hypotheses for how tectonic plates have moved over Earth’s history, reconstructions of the latitudinal biodiversity gradient (LBG) might also vary based on the GPM used. Here, using the fossil record of five major marine invertebrate groups, we evaluate the impact of GPM choice on reconstructions of the LBG over the Phanerozoic. Our results show that GPM choice can lead to different conclusions about the shape and strength of LBGs in deep time, even at a coarse spatial scale. These findings suggest additional caution is needed when reconstructing deep-time biogeographic patterns and macroevolutionary events, such as the origin of the present-day LBG. We therefore advocate for future palaeobiogeographic studies to conduct sensitivity analyses investigating the impact of GPM choice on their conclusions, and for greater interdisciplinary collaboration between palaeobiologists and palaeogeographic modellers to avoid common issues in the use of GPMs.

## Introduction

Today, species richness generally decreases from the tropics to the poles. This phenomenon, known as the ‘Latitudinal Biodiversity Gradient’, is one of Earth’s longest recognised and most prevalent macroecological patterns [1–4]. It has historically been observed across numerous taxonomic groups in the terrestrial [2,3,5] and marine realms [6,7], although there is evidence that it is currently experiencing a major transition, with diversity peaks moving away from the Equator [8–10]. The fossil record suggests that such changes in the shape and strength of latitudinal biodiversity gradients (LBGs) are to be expected, with flattened or bimodal gradients observed across a range of taxonomic groups at various points in Earth’s geological history [11–21]. Several studies have demonstrated that broadly unimodal gradients – with tropical peaks and poleward declines in taxonomic richness – have been restricted to the last 30 million years (Myr) and intervals of the Palaeozoic when cool icehouse climatic regimes dominated [4,16,21–24]. Conversely, during intervals of warmer climatic conditions (i.e. greenhouse and interglacial periods), some taxonomic groups are thought to have exhibited flattened LBGs, or even temperate peaks in biodiversity [12,13,15,17,18,25,26]. However, recent work suggests that our ability to recognise different types of LBGs in deep time is hindered by incomplete and heterogeneous spatial sampling in the fossil record, casting doubt on the reliability of these findings [27,28].

When examining macroecological patterns – such as the LBG – in the present day, neontologists use the geographic coordinates at which they collected their samples to infer the spatial distributions of their taxa. However, palaeobiologists must contend with the movement of tectonic plates over geological timescales, which results in the geographic coordinates at which a fossil was found not necessarily representing its location *in vivo*. Being able to accurately translate modern-day locality coordinates into the geographic distributions of fossil taxa is therefore a fundamental but difficult step in estimating the LBG in deep time. To conduct this translation, palaeobiologists routinely use Global Plate Models (GPMs), sometimes also referred to as ‘palaeorotation models’ or ‘plate rotation models’ [16,19,20,26,29,30]. These models constitute hypotheses for the tectonic evolution of the Earth, developed by modelling the motion of continental – and sometimes marine – plates across its surface through geological time. Many GPMs have been constructed [e.g. 31,32–34], varying in how they define the geological boundaries of continents, and the types of data which are used to validate them, having consequences for how fossil occurrences might be palaeogeographically reconstructed [35,36]. However, to date, few palaeobiological studies (though see refs. [37,38]) have considered how different GPMs might influence reconstructions of the latitudinal distributions of fossil occurrences.

Recent work has quantified the extent to which GPMs can differ in their estimation of the palaeogeographic distribution of fossil occurrences [36]. Here, we evaluate how sensitive deep-time reconstructions of the LBG are to GPM choice. Specifically, we test the extent to which the shape and strength of LBGs vary when using different GPMs, throughout the entire Phanerozoic (i.e. the last 540 Myr). To do so, we estimate the palaeogeographic distribution of fossil occurrences for five major marine invertebrate groups, using data from the Paleobiology Database [39], and four commonly applied GPMs. We then describe the raw latitudinal distribution of occurrences generated by each GPM, reconstruct the different LBGs using coverage-based rarefaction, and quantify the variability of these reconstructed gradients through time between GPMs.

## Materials and Methods

### Occurrence data

We downloaded Fortunian–Holocene (541–0 Ma) fossil occurrence data from the Paleobiology Database (PBDB; https://paleobiodb.org/) for five major marine invertebrate groups (Bivalvia, Brachiopoda, Cephalopoda, Gastropoda, Trilobita) on December 16 2024. Fossil occurrence data were downloaded using the PBDB Application Programming Interface (API) service and were restricted to marine environments and regular preservation (i.e. excluding form taxa and ichnotaxa). Occurrence data were subsequently binned into chrostratigraphic stage-level time bins following the International Geological Time Scale ver. 2023/09, with the exception of Holocene and Pleistocene stages which were collapsed into their equivalent Epoch-level bins (i.e. Holocene and Pleistocene) (Table S1). Temporal binning was carried out using the bin_time() function from the palaeoverse R package ver. 1.4.0, with the ‘majority’ approach [40]. As a result, all occurrences with less than 50% of their age range within their assigned geological stage were discarded. To minimise differences in the spatio-temporal resolution of the data, such as the lumping versus splitting of geological sections into PBDB collections, we rounded the geographic coordinates of all fossil collections to two decimal places and collapsed collections from the same time and place, removing any duplicated occurrences arising as a result. After data preparation, the occurrence dataset contained 344,258 fossil occurrences from 81,180 fossil (pseudo-) collections.

### Palaeogeographic reconstruction and palaeolatitudinal binning

To reconstruct the palaeogeographic distributions of fossil occurrences, we used their present-day coordinates and the midpoint age from their assigned geological stage with four GPMs: GOLONKA [32], TorsvikCocks2017 [41], PALEOMAP [33], and MERDITH2021 [34]. Different versions of the GOLONKA and PALEOMAP model have been extensively used throughout the geological and palaeontological literature [e.g. 15,16,17,19,38,42–44], and are the default models used in the Paleobiology Database [39]. In comparison, the TorsvikCocks2017 model has been less extensively used, while the more recent MERDITH2021 model is becoming increasingly applied [e.g. 37,45]. To generate palaeogeographic reconstructions we used the GPlates Web Service (https://gwsdoc.gplates.org) via the palaeorotate() function in palaeoverse ver. 1.4.0 [40]. For each GPM, we binned fossil occurrences into one of six equal-area latitudinal bins via the lat_bins_area() function in palaeoverse ver. 1.4.0 [40], using the estimated palaeolatitudes from the palaeogeographic reconstruction (Table S2). These latitudinal bins broadly represent three climatic zones within each hemisphere today: tropical, temperate, and polar. We recorded the number of fossil occurrences which were unable to be palaeogeographically reconstructed due to model incompatibility (e.g. fossil occurrence age estimate exceeding age of assigned plate) for each GPM.

### Quantifying the LBG

Subsequent data manipulation and visualisation were carried out using the Tidyverse suite of R packages [46] and the deeptime R package ver. 2.1.0 [47]. The raw number of genera were counted per stage per palaeolatitudinal bin for each GPM. We also estimated the LBG using coverage-based interpolation and extrapolation of Hill numbers with a coverage level of 0.4, using the R package iNEXT ver. 3.0.1 [48]. The rarefaction portion of this approach is equivalent to the method commonly known as Shareholder Quorum Subsampling (SQS) amongst palaeontologists [49,50]. Extrapolated values with an estimated sample size more than double that of the observed sample size were discarded, as recommended by ref [48]. As we were principally interested in the shape and strength of the LBG, all reconstructed gradients were normalised by dividing the value for each latitudinal bin by that of the latitudinal bin with peak richness within each stage.

We compared estimated LBGs using two approaches. Firstly, for each stage and hemisphere, we determined the palaeolatitudinal bin with maximum estimated richness for each GPM. We then calculated the number of stages in which estimated gradients agree on the spatial bin with peak diversity, as well as the number of bins with a low, mid-, and high palaeolatitudinal peak in diversity. Secondly, we calculated the mean pairwise rank order difference between GPMs. That is, for each stage and GPM, the palaeolatitudinal bins were put in rank order of richness, and the differences between models were computed and the mean calculated. To enable fair comparison between stages with differing numbers of occupied spatial bins, average rank order differences were normalised by maximum possible average difference given the number of ranks available. The former of these metrics tests whether models agree on where peak diversity is concentrated in each GPM, while the latter tests for differences in the estimated ordered distribution of diversity (e.g. low-to-mid-to-high, mid-to-low-to-high, etc.).

As the estimated latitudinal distribution of biodiversity is closely tied to the distribution of fossil localities, we also computed the number of fossil localities in each palaeolatitudinal bin and stratigraphic stage for each GPM. Furthermore, to test whether reconstructions of the LBG are more sensitive to GPM choice with increasing age of rotation, we calculated the median pairwise palaeolatitudinal difference and great-circle distance (otherwise known as the Haversine distance) between GPMs for each palaeogeographically reconstructed fossil collection. As we were only interested in palaeolatitudinal differences between models given the nature of our study, we set palaeolongitudinal coordinates to 0º during calculations of great-circle distance (i.e. we calculate the palaeolatitudinal great-circle distance). We include this metric in addition to palaeolatitudinal differences–measured in degrees of latitude–to provide a standard measure of geographic distance.

## Results

### Palaeogeographic reconstructions

Palaeogeographic reconstruction of fossil occurrences reveals that the number of fossil localities (unique spatiotemporal data points, *n* = 47180) available for allocation to palaeolatitudinal bins – localities which palaeocoordinates could be generated for – varies with the GPM used (Fig. 1). While in most geological stages, the number of localities available for allocation to palaeolatitudinal bins is similar, it varies considerably for others (Fig. 1). For example, of the 601 fossil localities assigned to the Lutetian (Cenozoic), palaeocoordinates could be generated for 597 (∼99%) fossil localities when using the PALEOMAP model, 592 (∼99%) with GOLONKA, 546 (∼90%) with MERDITH2021, and 496 (∼83%)with TorsvikCocks2017. Similarly, of the 879 fossil localities available for the Anisian (Triassic), palaeocoordinates were generated for 853 (∼97%), 837 (∼95%), and 771 (∼88%) using the GOLONKA, MERDITH2021, and PALEOMAP models, respectively. However, just 553 (∼63%) fossil localities could be reconstructed when using the TorsvikCocks2017 model. This, along with spatial discrepancies in palaeogeographic reconstructions between GPMs, resulted in the number of samples allocated to each palaeolatitudinal bin (low, medium, high palaeolatitudes) varying with GPM (Figure S1). Of the 97 geological stages, the number of palaeolatitudinal bins containing enough samples (i.e. fossil collections) for estimating genus richness also varies between GPMs, with a difference of three bins for 1 geological stage (∼1%), two bins for 17 geological stages (∼18%), one bin for 47 stages (∼48%), and zero bins for 32 stages (∼33%) (Fig. 1).

**Figure 1.**
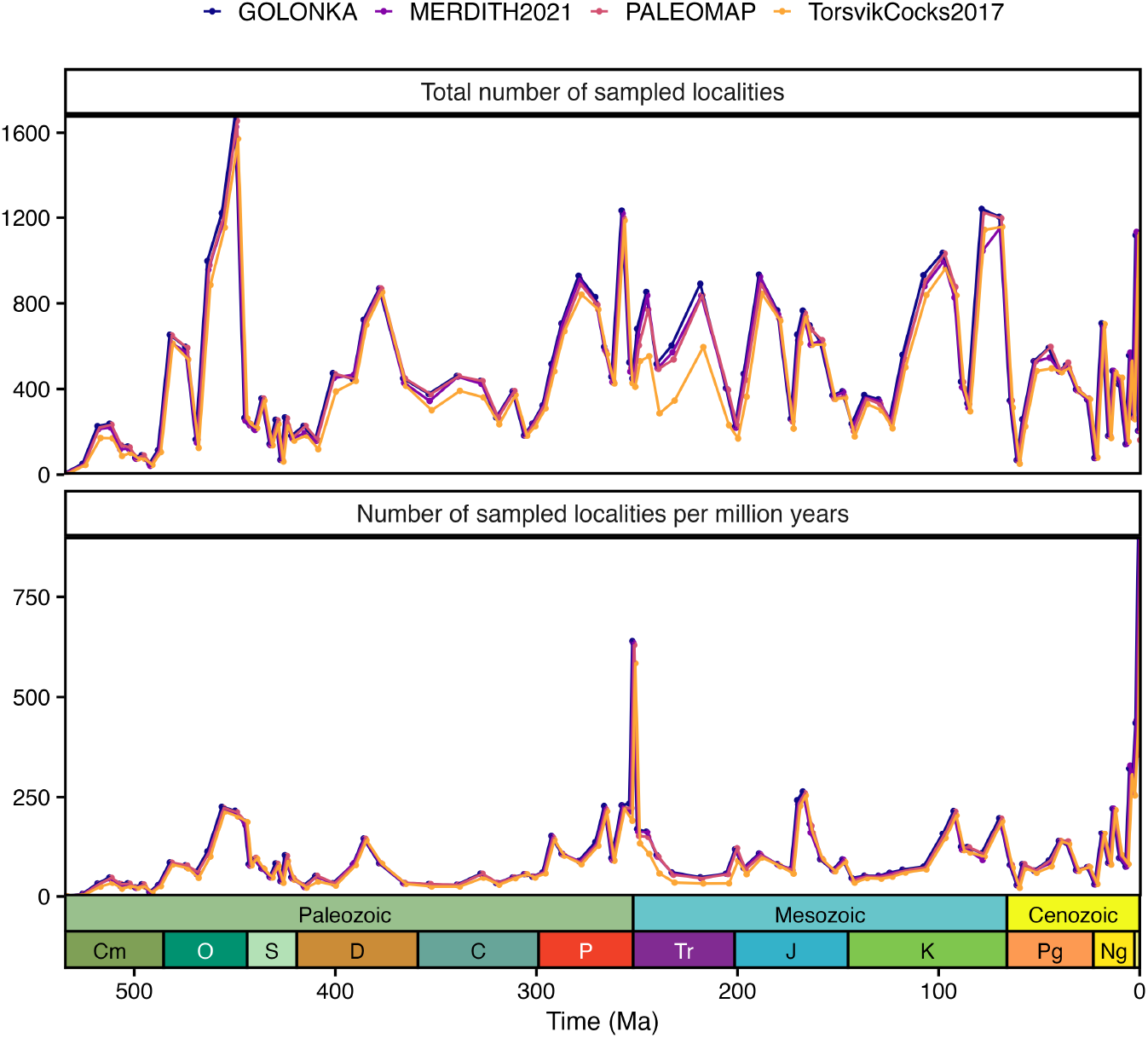
Number of fossil localities within each Phanerozoic stage that could be palaeogeographically reconstructed using four Global Plate Models: GOLONKA [32], PALEOMAP [33], MERDITH2021 [34], and TorsvikCocks2017 [41]. (a) Raw counts of the number of fossil localities available for palaeolatitudinal binning per stage from each Global Plate Model. (b) The number of localities per million years available for palaeolatitudinal binning per stage from each Global Plate Model. Period abbreviations are as follows: Cambrian (Cm); Ordovician (O), Silurian (S), Devonian (D), Carboniferous (C), Permian (P), Triassic (Tr), Jurassic (J), Cretaceous (K), Paleogene (Pg) and Neogene (Ng). The Quaternary is not depicted. The geological time scale axis was added to the plot using the R package ‘deeptime’ ver. 2.1.0 [47]. Note, the plotted data has been slightly dodged along the x-axis to improve readability.

Analysis of the average pairwise palaeolatitudinal difference and pairwise great-circle distance between palaeogeographically reconstructed fossil collections indicates, on average, an increasing difference between GPMs with age of reconstruction (Fig. 2). Specifically, average median pairwise palaeolatitudinal differences between models for each stage is 2.51º latitude for the Cenozoic, 4.49º latitude for the Mesozoic, and 10.34º latitude for the Palaeozoic. Similarly, average median pairwise great-circle distance between models for each stage is ∼254 km for the Cenozoic, ∼477 km for the Mesozoic, and ∼1093 km for the Palaeozoic. However, while there is a general increasing difference between the palaeogeographic reconstruction of fossil collections with age of reconstruction, there are intervals of low palaeolatitudinal differences, for example the Permian (Fig. 2). Nevertheless, prior to the Late Jurassic, there is generally a larger variability (spread) in palaeolatitudinal difference and great-circle distance than during the Late Jurassic, Cretaceous and Cenozoic (Fig. 2). For example, palaeolatitudinal differences can be larger than 25º latitude for some reconstructed fossil collections of the Jurassic (*n* = 488), Triassic (*n* = 322), Permian (*n* = 222), Carboniferous (*n* = 103), Devonian (*n* = 155), Silurian (*n* = 46), Ordovician (*n* = 680), and Cambrian (*n* = 847).

**Figure 2.**
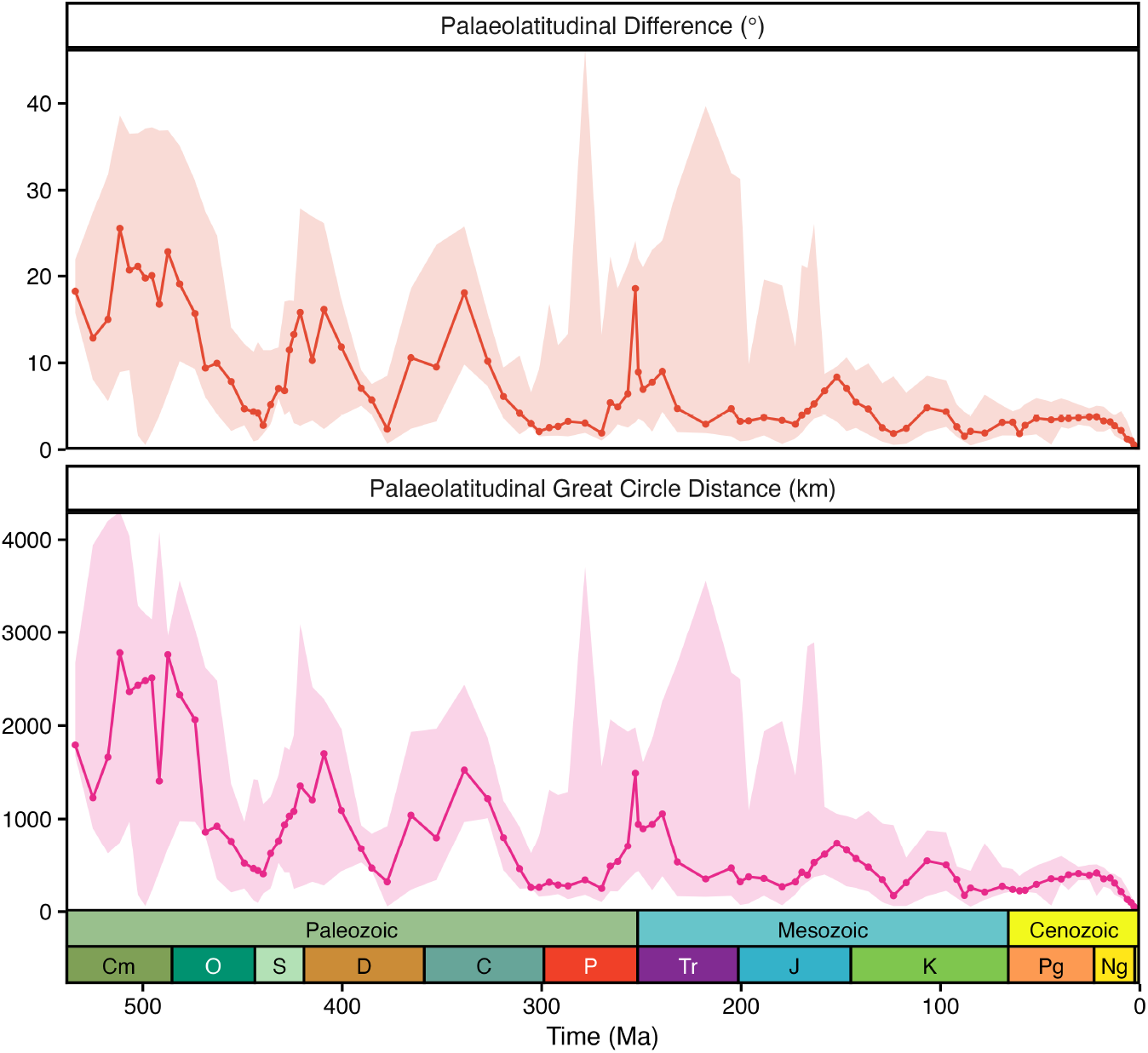
Phanerozoic trends in the spatial discrepancies between palaeogeographic reconstructions of fossil collections for four Global Plate Models: GOLONKA [32], PALEOMAP [33], MERDITH2021 [34], and TorsvikCocks2017 [41]. (a) Average (median; 5th and 95th percentiles) pairwise palaeolatitudinal distance between palaeogeographic reconstructions for fossil collections. (b) Average (median; 5th and 95th percentiles) pairwise great-circle distance between palaeogeographic reconstructions for fossil collections. In both panels, the ribbon depicts the 5th and 95th percentiles of the data. Period abbreviations are as follows: Cambrian (Cm); Ordovician (O), Silurian (S), Devonian (D), Carboniferous (C), Permian (P), Triassic (Tr), Jurassic (J), Cretaceous (K), Paleogene (Pg) and Neogene (Ng). The Quaternary is not depicted. The geological time scale axis was added to the plot using the R package ‘deeptime’ ver. 2.1.0 (Gearty, 2023).

The shapes of the reconstructed LBGs for each geological stage are mostly consistent between GPMs (Fig. 3; Fig. S2). For all four GPMs, we observe the same overall trend in LBGs over geological time, namely that the peak in diversity tended to lie in the Southern Hemisphere prior to the Permian, but shifted to the Northern Hemisphere after the Permian, in all four models (Fig. 4; Fig. S3). Throughout the Phanerozoic, peak diversity tended to appear at low and mid-latitudes (Fig. 4; Table S3; Fig. S3). However, during some intervals, we see considerable variation in the shape of the LBG between GPMs, for example during the Guzhangian, Tremadocian, Sakmarian, Induan, and Oxfordian (Fig. 3; Fig. S2). Notably, these differences are not restricted to older intervals (Fig. S4–5), with some intervals of the Cenozoic (e.g. Serravallian, Burdigalian) demonstrating opposing trends in the palaeolatitudinal distribution of biodiversity (Fig. 3; Fig. S2).

**Figure 3.**
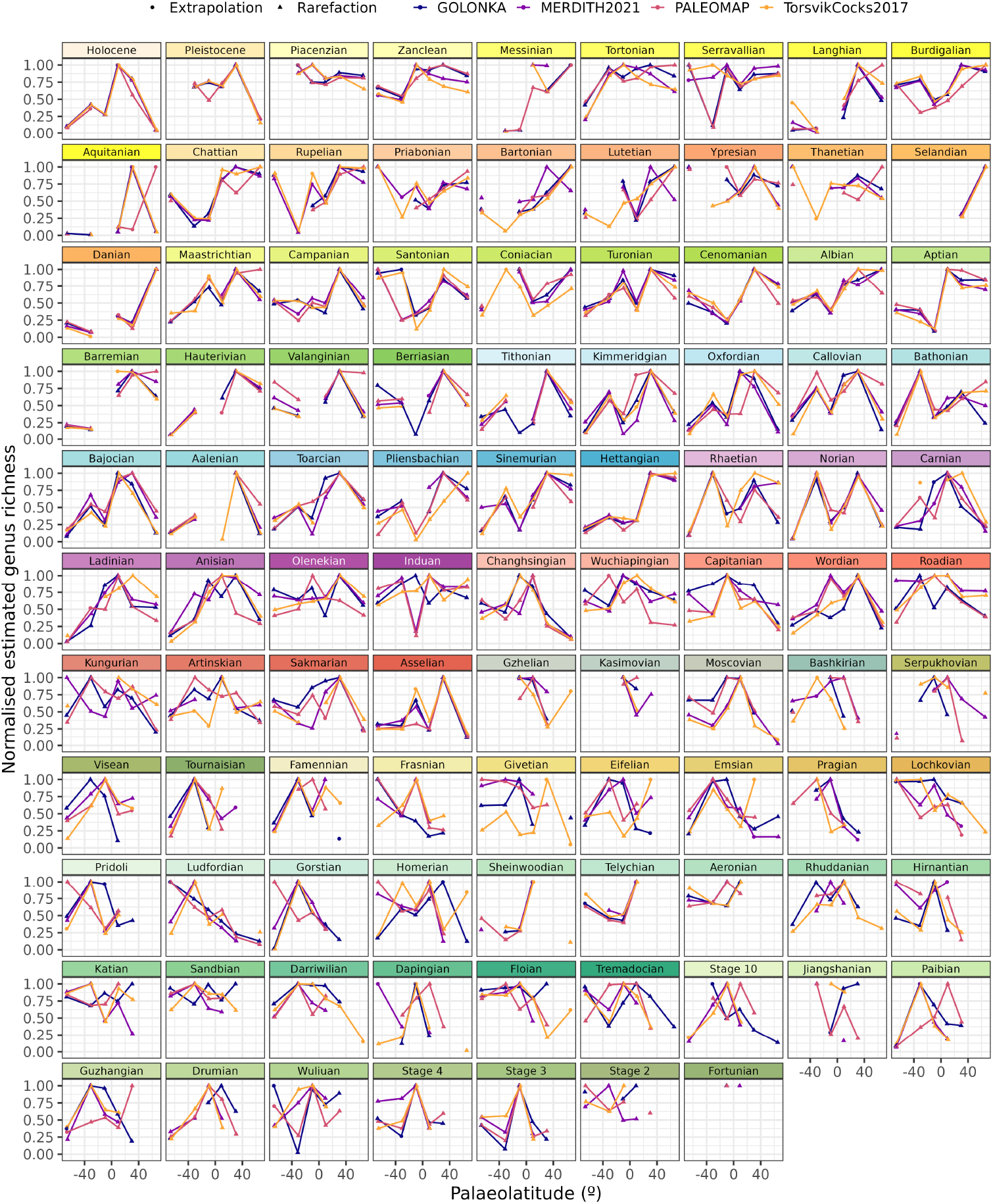
Phanerozoic stage-level reconstructions of the latitudinal biodiversity gradient for five major marine invertebrate groups (Bivalvia, Brachiopoda, Cephalopoda, Gastropoda, and Trilobita). Each individual plot depicts the estimated normalised genus richness within each palaeolatitudinal bin for four Global Plate Models (GPMs): GOLONKA [32], PALEOMAP [33], MERDITH2021 [34], and TorsvikCocks2017 [41]. Genus richness was estimated for each stage, palaeolatitudinal bin, and GPM using the iNEXT R package ver. 3.0.1 [48] with a coverage level (otherwise known as a quorum level) of 0.4. Genus richness was normalised for each stage and GPM by dividing the genus richness within each palaeolatitudinal bin by the maximum value across palaeolatitudinal bins. Note, the plotted data has been slightly dodged along the x-axis to improve readability.

**Figure 4.**
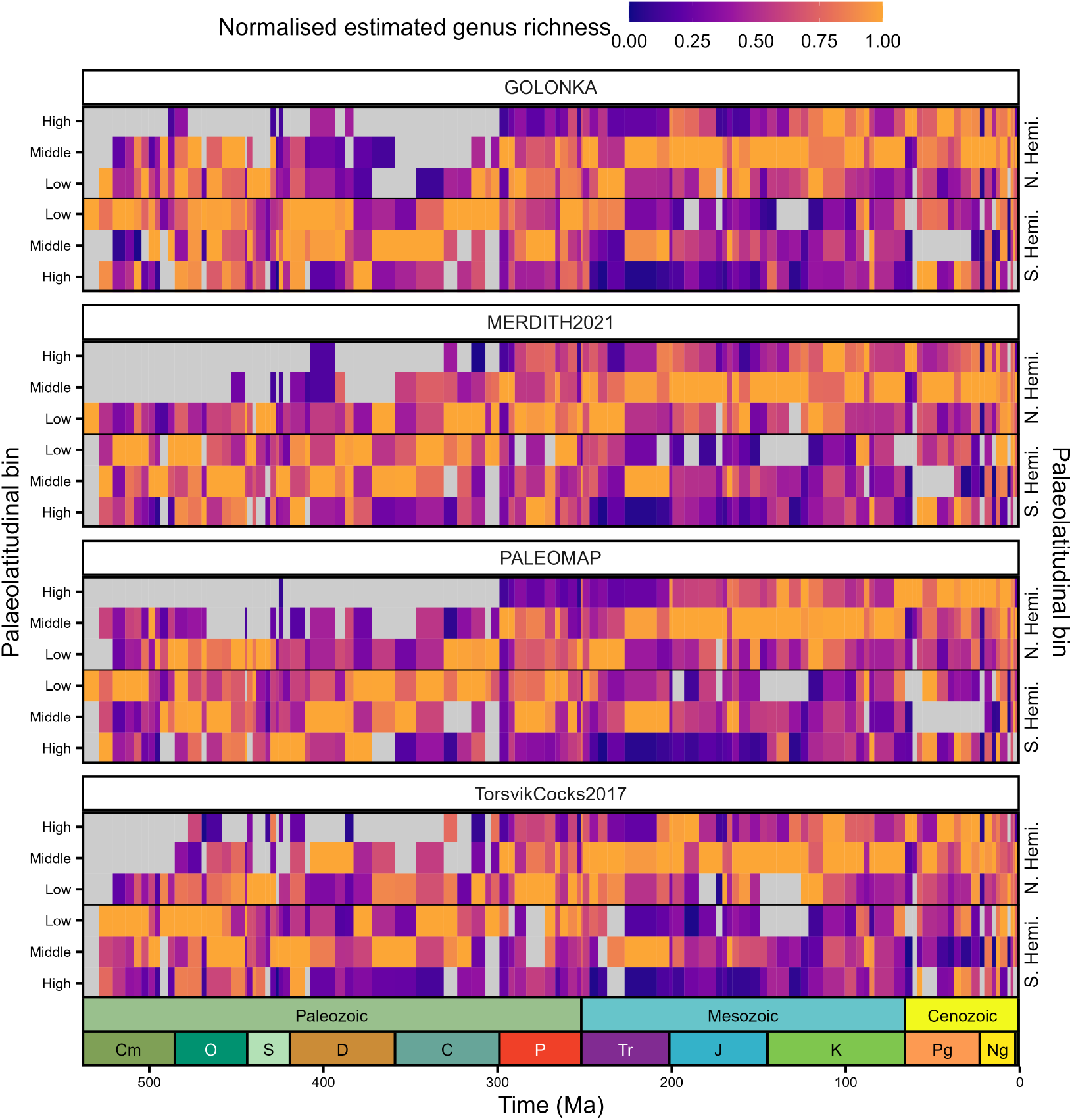
Phanerozoic heatmaps of normalised estimated genus richness for five major marine invertebrate groups (Bivalvia, Brachiopoda, Cephalopoda, Gastropoda, and Trilobita). Each individual panel depicts the normalised estimated genus richness (coverage-based rarefaction) for four Global Plate Models (GPMs): GOLONKA [32], PALEOMAP [33], MERDITH2021 [34], and TorsvikCocks2017 [41]. Heatmaps depict the normalised estimated genus richness within palaeolatitudinal bins (low, middle, and high) within the Northern and Southern Hemisphere. Genus richness was estimated for each stage, palaeolatitudinal bin, and GPM using the iNEXT R package ver. 3.0.1 [48] with a coverage level (otherwise known as a quorum level) of 0.4. Genus richness was normalised for each stage and GPM by dividing the genus richness within each palaeolatitudinal bin by the maximum value across palaeolatitudinal bins. Grey tiles indicate palaeolatitudinal bins without sufficient data to generate richness estimates.

We quantified the differences between diversity estimates obtained using the four GPMs by evaluating the extent to which the models agreed on which palaeolatitudinal bin contained the most diversity in each hemisphere (Fig. 5; Fig. S6). Our results show that the extent to which peaks in reconstructed palaeolatitudinal diversity were consistent between models was variable through time, with only ∼19% of stages in the Palaeozoic, ∼47% in the Mesozoic, and ∼26% in the Cenozoic showing agreement between all four models. Notably, this marginally differed between the two hemispheres, with ∼49% agreement in the Northern Hemisphere and ∼43% in the Southern Hemisphere over the whole Phanerozoic (Fig. 5). We also compared the shape of the LBGs by calculating the normalised average rank order difference of palaeolatitudinal bins between each pair of models. Overall, there was most agreement between the models on the rank order of richness in palaeolatitudinal bins during the Mesozoic; no pair of models disagreed by more than ∼0.78, with a mean value of ∼0.25 (Fig. 6). However, during the Palaeozoic (mean = ∼0.45) and Cenozoic (mean = ∼0.27), in some cases, pairs of models wholly disagreed on the rank order of diversity in palaeolatitudinal bins (normalised rank order difference of 1). This was less prevalent when calculated using the raw richness compared to the coverage-based diversity estimates (Fig. S7), likely due to the lower sample size threshold for the former, meaning a higher number of spatial bins contained enough data to be analysed.

**Figure 5.**
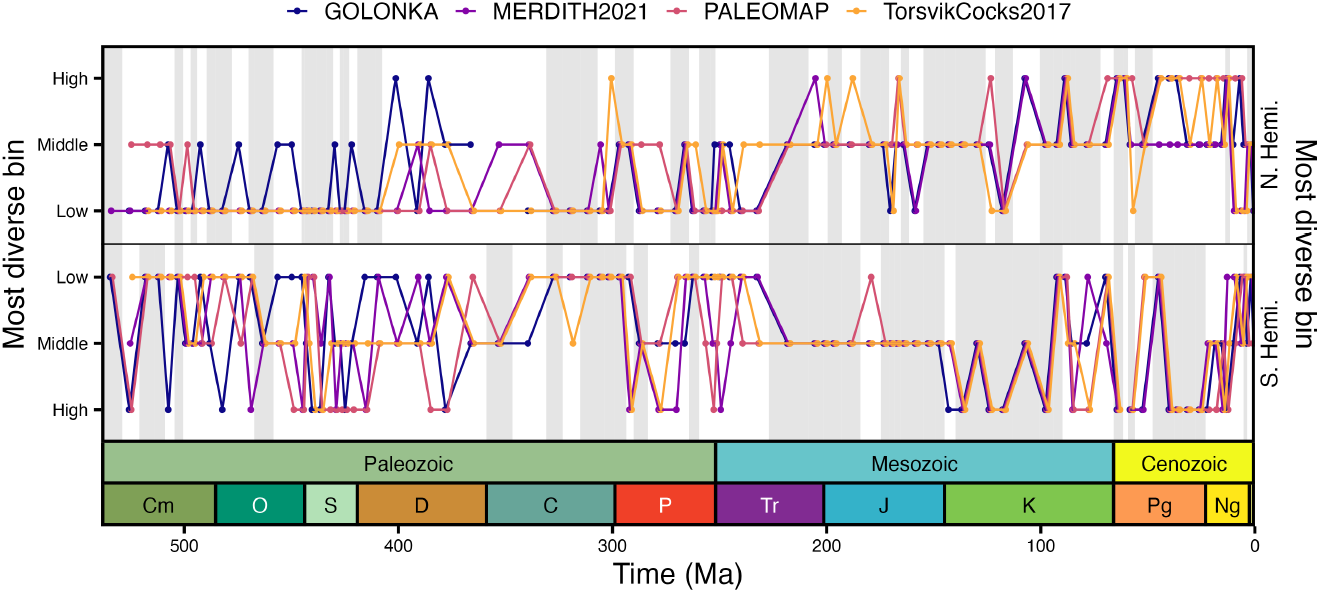
Phanerozoic plot of the palaeolatitudinal bin (low, middle, high) with the greatest estimated genus richness within the Northern Hemisphere and Southern Hemisphere for each Global Plate Model (GPM): GOLONKA [32], PALEOMAP [33], MERDITH2021 [34], and TorsvikCocks2017 [41]. Grey shading highlights intervals in which the palaeolatitudinal bin with the greatest estimated genus richness is in agreement across all GPMs. Genus richness was estimated for each stage, palaeolatitudinal bin, and GPM using the iNEXT R package ver. 3.0.1 [48] with a coverage level (otherwise known as a quorum level) of 0.4. Period abbreviations are as follows: Cambrian (Cm); Ordovician (O), Silurian (S), Devonian (D), Carboniferous (C), Permian (P), Triassic (Tr), Jurassic (J), Cretaceous (K), Paleogene (Pg) and Neogene (Ng). The Quaternary is not depicted. The geological time scale axis was added to the plot using the R package ‘deeptime’ ver. 2.1.0 [47]. Note, the plotted data has been slightly dodged along the x-axis to improve readability.

**Figure 6.**
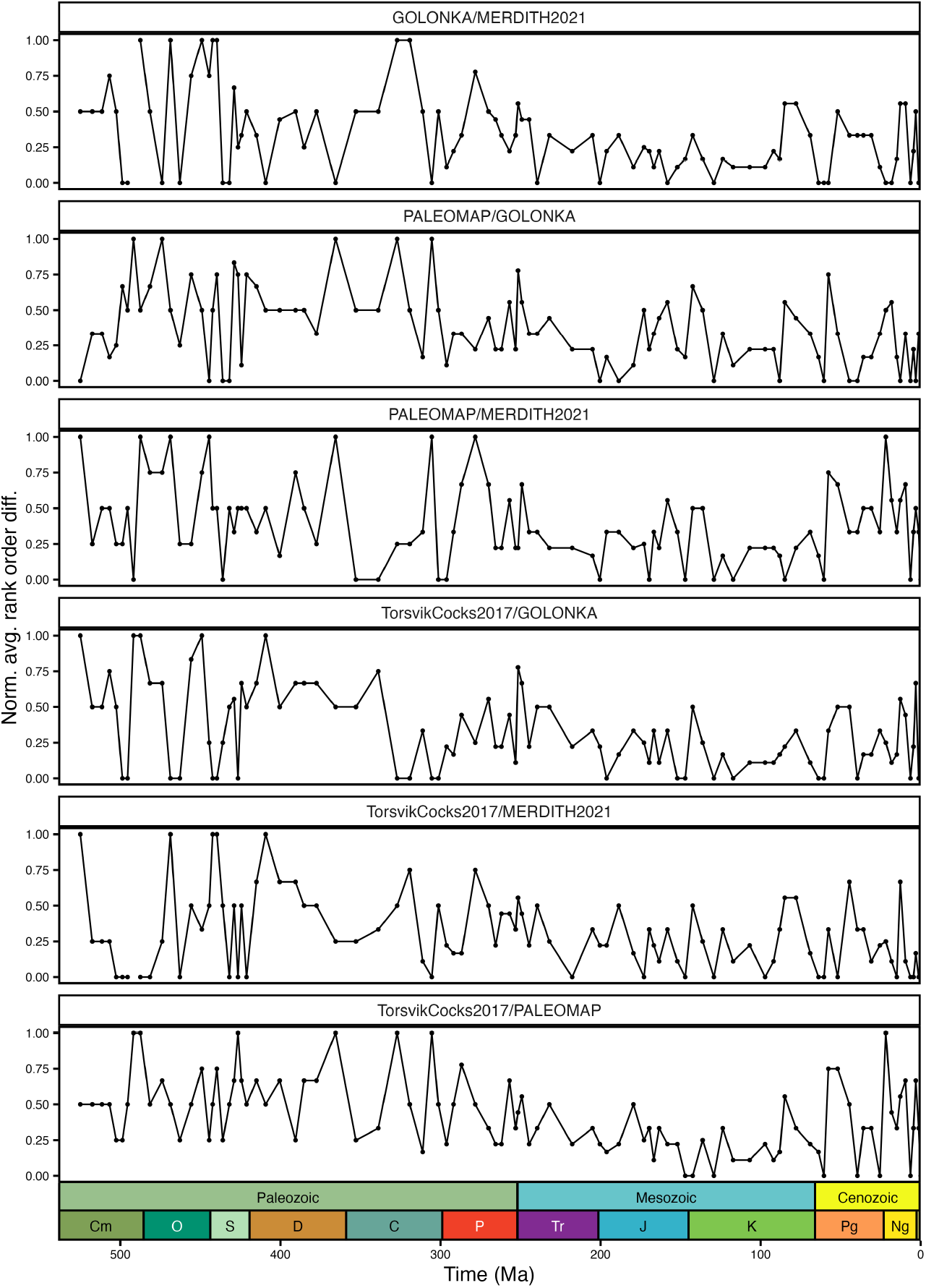
Phanerozoic plot of the normalised average rank order pairwise difference between latitudinal biodiversity gradients (estimated genus richness) for each Global Plate Models (GPMs): GOLONKA [32], PALEOMAP [33], MERDITH2021 [34], and TorsvikCocks2017 [41]. Genus richness was estimated for each stage, palaeolatitudinal bin, and GPM using the iNEXT R package ver. 3.0.1 [48] with a coverage level (otherwise known as a quorum level) of 0.4. Average rank order differences are normalised by maximum possible average difference given the number of ranks available. Period abbreviations are as follows: Cambrian (Cm); Ordovician (O), Silurian (S), Devonian (D), Carboniferous (C), Permian (P), Triassic (Tr), Jurassic (J), Cretaceous (K), Paleogene (Pg) and Neogene (Ng). The Quaternary is not depicted. The geological time scale axis was added to the plot using the R package ‘deeptime’ ver. 2.1.0 [47].

## Discussion

Global Plate Models (GPMs) are regularly used in palaeobiology to reconstruct the palaeogeographic distribution of fossil occurrences e.g. refs [16,17,19,20,26,29,30,37]. In this study, we evaluated whether GPM choice impacts reconstruction of the latitudinal biodiversity gradient (LBG) over the Phanerozoic. This gradient is one of the most universal patterns in macroecology today [2,3], and its accurate characterisation is dependent on the latitude at which occurrences are believed to have been observed. Our results demonstrate that choice of GPM when estimating the palaeolatitude of fossil occurrences (latitudinal position at time of deposition) can have a strong influence on the reconstructed LBG, including when using simple metrics such as the palaeolatitudinal bin with peak biodiversity (Fig. 5).

Previous work using the fossil record suggests that the LBG has varied in shape and strength through time, with previous analyses demonstrating unimodal, flattened, and bimodal gradients at various points in Earth’s geological history [e.g. 11,12–15,17–21]. Our results broadly agree with these findings, indicating that the strength and shape of the LBG has varied through time, regardless of which GPM is used to reconstruct the palaeogeographic distribution of fossil occurrences. Previous work has also shown that apparent tropical peaks and poleward declines in taxonomic richness appear to be restricted to the last 30 million years, and intervals of the Palaeozoic when cool icehouse climatic regimes persisted [e.g. 4,20,21,23]. Conversely, during intervals of warmer climatic conditions (i.e. greenhouse and interglacial periods), various taxonomic groups have exhibited flattened LBGs, or even temperate peaks in biodiversity [e.g. 11,14,15,17,18,20]. These studies suggest that climate is a major driver of the LBG over evolutionary timescales. However, the extent to which these observed latitudinal patterns are the result of inherent biases (e.g. variable fossil preservation, stratigraphic completeness), as opposed to biological processes, remains uncertain [e.g. 27,28,51]. Although our results support the notion of a dynamic gradient through time, we demonstrate that the strength and the shape of the LBG can vary depending on the GPM selected for palaeogeographic reconstruction. While this is most apparent in older geological stages, such as those of the Palaeozoic (Fig. 3; Fig. 5; Fig. 6), this can also impact reconstructions in more recent intervals such as the Lutetian, in the Cenozoic (Fig. 3). However, in more recent intervals, differences between LBG reconstructions are a result of differences in the number of fossil localities for which palaeogeographic coordinates can be generated with each GPM, rather than actual palaeogeographic differences between them. This is principally driven by differences in the described motion of plates (or timing of tectonic events) through time for different GPMs. For example, in one GPM, an area of a tectonic plate may originate earlier than in another, and if the age estimate of a fossil occurrence exceeds the age range of the assigned plate, no palaeogeographic coordinates can be generated. For example, of the 52 Tortonian fossil localities from Panama, palaeogeographic coordinates could be generated for every locality using the PALEOMAP and TorsvikCocks2017 models. However, no palaeogeographic coordinates could be generated for these localities using the GOLONKA and MERDITH2021 models.

Although during most geological stages there is broad agreement in the shape of the LBG between GPMs (Fig. 3), there are some for which this is not the case. For example, in 25 stage-hemisphere combinations (7 Northern Hemisphere, 18 Southern Hemisphere), disagreement between models can be as extreme as concluding that a LBG has a tropical peak in biodiversity under one model, and a polar peak in biodiversity under another, such as in the Southern Hemisphere during the Givetian (Fig. 3; Fig. 5). As such, the magnitude of palaeolatitudinal uncertainty represented across different GPMs can lead to markedly different LBG reconstructions and subsequent interpretations, even at a coarse spatial scale. This is most extreme in the Southern Hemisphere, likely a consequence of the lower number of occurrences and reduced sampling coverage available for this hemisphere, resulting in a higher sensitivity to small differences in the data. Nevertheless, in less extreme examples, 68 stage-hemisphere combinations (33 Northern Hemisphere, 35 Southern Hemisphere) still demonstrate that the LBG would be reconstructed as having a tropical peak in diversity under one GPM, and a temperate peak in diversity under another. Consequently, in order to robustly test whether certain types of gradients are limited to specific climate states (e.g. unimodal gradients in icehouse periods), consideration of a range of GPMs is required. Across temporal scales, differences in the reconstruction of LBGs between GPMs could also impact conclusions about the timing of events such as the origin of the present-day LBG [e.g. 21]. This is essential in enabling us to test the evolutionary drivers of such events, for which their timing must be robustly constrained.

Using extrapolation as a means to address sampling bias may also be contributing to the strength of contrasts between GPMs: our approach uses relative abundance data to determine how much extrapolation is required, meaning that estimates of richness can be highly sensitive to the shifting of a handful of collections between palaeolatitudinal bins. The LBGs reconstructed using the raw data are more consistent with each other than those produced using coverage-based extrapolation and rarefaction (Fig. 3; Fig. S2). This could support the notion that our extrapolation approach is overly sensitive, but may also be due to a stronger signal of sampling bias pushing the raw LBGs closer to each other.

The reconstruction of regional palaeobiodiversity is complicated. Despite the abundance of data within the Paleobiology Database, there remain numerous gaps in our dataset, illustrated by the absence of data in many individual palaeolatitudinal bins and even in a small number of entire temporal stages (Fig. 3; Fig. S2). While some of these gaps are attributable to a lack of shallow marine shelf area within certain spatio-temporal bins, many might be filled through additional fossil data collection in the future. Ideally, we would have preferred to subdivide data into more than six latitudinal bins for analyses. However, unfortunately, higher spatial resolution in this case results in noisier and patchier latitudinal gradients which are governed more strongly by sampling bias than by true diversity trends. Furthermore, while many recent spatial palaeobiodiversity studies have relied on using equal-area rectangular or hexagonal cells (e.g. refs [19,52,53]), further subdivision of our latitudinal bins would likewise introduce more noise into our analyses.

Another source of potential limitation lies in how we quantified the LBGs. We decided to quantify the most diverse bin for each model and calculate the average rank order difference between pairwise models. Both of these metrics have the benefit of utilising relative diversity rather than raw differences, which should be less susceptible to sampling biases, especially when coverage-based rarefaction is used. We found the rank order difference to be particularly promising as a way to quantify the relative differences between two gradients. However, the maximum average rank order difference is highly dependent on the number of bins that are sampled. We normalised for this dependence; however, it is important to note that with fewer bins it takes fewer rank order changes to achieve a higher normalised score.

There exist many other ways to quantify LBGs. For example, the sum of squares could be used to calculate the raw deviation of one gradient from another. However, such a calculation could be heavily biased by sampling intensity. Another option would be to compare the fit of various regression models to assess the modality and shape of the gradients [19]. However, the sensitivity and success of this type of approach are heavily dependent on the number of latitudinal bins available for fitting. Finally, a variety of null models, such as the modern unimodal gradient or the bimodal gradient of past greenhouse periods [4], could be established. The deviation of an observed gradient from these null models could be used to assess how well the gradient is explained by a particular hypothesis. However, this approach has its own set of limitations, including how the set of null models/hypotheses is selected and where the null gradients are centred latitudinally. The assessment of these various alternative approaches is outside of the scope of this study. However, we believe that any of these other methods would echo our findings that GPM choice can be critical in reconstructing LBGs in deep time.

Determining the palaeocoordinates at which organisms’ remains were deposited is a fundamental step in many macroecological and macroevolutionary analyses, and although our results indicate that caution is necessary in doing this, we have additional, specific recommendations. Firstly, we recommend that researchers carefully consider which GPM is most suitable for their work. The spatiotemporal coverage of different GPMs, the proportion of occurrence data for which palaeogeographic coordinates can be generated, the data types taken into account when producing the models, and which models were previously used for similar analyses, might all be relevant to making an informed decision. Secondly, we strongly advise consideration of palaeogeographic uncertainty between different GPMs, along with the evaluation of the potential impact of model choice on conclusions. For example, workflows could be implemented using multiple different GPMs, with palaeogeographic uncertainty included in figures through error bars, or results generated with different GPMs included in supplementary material. Finally, we recommend greater collaboration between palaeobiologists and palaeogeographic modellers in future palaeobiogeographic studies [e.g. 36]. Common issues such as the use of GPMs in the mantle reference frame, and confusion over nomenclature, may be reduced through greater exchange between those building GPMs and those using them in secondary research [35].

## Conclusion

Our study demonstrates that reconstructing deep time LBGs is not a trivial task. The use of one Global Plate Model over another can lead to different reconstructions of the palaeolatitudinal distribution of biodiversity, influencing conclusions about the shape and strength of latitudinal biodiversity gradients in deep time, as well as constraining macroevolutionary events such as the origin of the present-day latitudinal biodiversity gradient. In general, we find support for our hypothesis which states reconstructions of the latitudinal biodiversity gradient are more sensitive to Global Plate Model choice with increasing age of rotation. Specifically, our findings show that prior to the Mesozoic, differences between Global Plate Models become increasingly larger with age of rotation. However, we also find that even small differences in the palaeogeographic reconstruction of fossil occurrences between models can have a large impact on reconstructions of the latitudinal biodiversity gradient in younger intervals (e.g. Aquitanian, Burdigalian, Langhian, Tortonian and Messinian). For future palaeobiogeographic studies, we recommend greater interdisciplinary collaboration between palaeobiologists and palaeogeographic modellers to avoid common issues in the use of Global Plate Models. We also endorse that future studies test the sensitivity of their conclusions to Global Plate Model choice, and quantify the palaeogeographic uncertainty associated with their data.

## Data accessibility

The data and code generated in this study have been included within the paper, its supplementary material, and dedicated GitHub repository: https://github.com/LewisAJones/LBG-GPM-choice. In addition, they have been uploaded to a dedicated Zenodo repository: https://doi.org/10.5281/zenodo.14616817.

## Authors’ contributions

L.A.J conceived the project. All authors contributed to the development of the project. All authors contributed to the writing of the manuscript. All authors contributed to data analyses. W.G. and L.B. produced the figures.

## Acknowledgements

We are grateful for the efforts of all those who have collected and entered data into the Paleobiology Database. The contributions of L.A.J. were supported by a UK Natural Environment Research Council (NERC) Independent Research Fellowship and a Norman Newell Early Career Grant from the Paleontological Society. The contributions of B.J.A. were supported by ETH Zürich. The contributions of W.G. were supported by the Lerner-Gray Postdoctoral Research Fellowship from the Richard Gilder Graduate School at the American Museum of Natural History and a Norman Newell Early Career Grant from the Paleontological Society. The contributions of L.B. were supported by a grant from the École Normale Supérieure de Lyon. This is Paleobiology Database official publication number XXX.

## Notes

https://github.com/LewisAJones/LBG-GPM-choice

https://doi.org/10.5281/zenodo.14616817

### Competing Interest Statement

The authors have declared no competing interest.

