## Supplementary Material 1 for "Global plate model choice impacts reconstructions of the latitudinal biodiversity gradient"

—

Lewis A. Jones[1], William Gearty[2,3], Lucas Buffan[4], and Bethany J. Allen[5,6]

—

[1]Department of Earth Sciences, University College London, Gower Street, London, WC1E 6BT, UK

[2]Division of Paleontology, American Museum of Natural History, New York, NY, 10024, USA

[3]Open Source Program Office, Syracuse University, Syracuse, NY, 13244, USA

[4]Institut des Sciences de l'Évolution de Montpellier, Université de Montpellier, CNRS, IRD, Place Eugène Bataillon, 34095 Montpellier cedex 05, France

[5]Department of Biosystems Science and Engineering, ETH Zürich, 4056 Basel, Switzerland

[6]Computational Evolution Group, Swiss Institute of Bioinformatics, 1015 Lausanne, Switzerland

—

#### Supplementary tables

**Table S1:** Chronostratigraphic stage-level time bins used in this study. Time bins follow the International Geological Timescale Ver. 2023/09 with the exception of the Holocene and Pleistocene stages, which have been collapsed into their equivalent Epoch-level bins.

| Bin | Interval | Maximum Age (Ma) | Minimum Age (Ma) | Duration (Myr) |
|---|---|---|---|---|
| 1 | Fortunian | 538.8000 | 529.0000 | 9.8000 |
| 2 | Stage 2 | 529.0000 | 521.0000 | 8.0000 |
| 3 | Stage 3 | 521.0000 | 514.0000 | 7.0000 |
| 4 | Stage 4 | 514.0000 | 509.0000 | 5.0000 |
| 5 | Wuliuan | 509.0000 | 504.5000 | 4.5000 |
| 6 | Drumian | 504.5000 | 500.5000 | 4.0000 |
| 7 | Guzhangian | 500.5000 | 497.0000 | 3.5000 |
| 8 | Paibian | 497.0000 | 494.0000 | 3.0000 |
| 9 | Jiangshanian | 494.0000 | 489.5000 | 4.5000 |
| 10 | Stage 10 | 489.5000 | 485.4000 | 4.1000 |
| 11 | Tremadocian | 485.4000 | 477.7000 | 7.7000 |
| 12 | Floian | 477.7000 | 470.0000 | 7.7000 |
| 13 | Dapingian | 470.0000 | 467.3000 | 2.7000 |
| 14 | Darriwilian | 467.3000 | 458.4000 | 8.9000 |
| 15 | Sandbian | 458.4000 | 453.0000 | 5.4000 |
| 16 | Katian | 453.0000 | 445.2000 | 7.8000 |
| 17 | Hirnantian | 445.2000 | 443.8000 | 1.4000 |
| 18 | Rhuddanian | 443.8000 | 440.8000 | 3.0000 |
| 19 | Aeronian | 440.8000 | 438.5000 | 2.3000 |
| 20 | Telychian | 438.5000 | 433.4000 | 5.1000 |
| 21 | Sheinwoodian | 433.4000 | 430.5000 | 2.9000 |
| 22 | Homerian | 430.5000 | 427.4000 | 3.1000 |
| 23 | Gorstian | 427.4000 | 425.6000 | 1.8000 |

| | | | | |
|---|---|---|---|---|
| 24 | Ludfordian | 425.6000 | 423.0000 | 2.6000 |
| 25 | Pridoli | 423.0000 | 419.2000 | 3.8000 |
| 26 | Lochkovian | 419.2000 | 410.8000 | 8.4000 |
| 27 | Pragian | 410.8000 | 407.6000 | 3.2000 |
| 28 | Emsian | 407.6000 | 393.3000 | 14.3000 |
| 29 | Eifelian | 393.3000 | 387.7000 | 5.6000 |
| 30 | Givetian | 387.7000 | 382.7000 | 5.0000 |
| 31 | Frasnian | 382.7000 | 372.2000 | 10.5000 |
| 32 | Famennian | 372.2000 | 358.9000 | 13.3000 |
| 33 | Tournaisian | 358.9000 | 346.7000 | 12.2000 |
| 34 | Visean | 346.7000 | 330.9000 | 15.8000 |
| 35 | Serpukhovian | 330.9000 | 323.2000 | 7.7000 |
| 36 | Bashkirian | 323.2000 | 315.2000 | 8.0000 |
| 37 | Moscovian | 315.2000 | 307.0000 | 8.2000 |
| 38 | Kasimovian | 307.0000 | 303.7000 | 3.3000 |
| 39 | Gzhelian | 303.7000 | 298.9000 | 4.8000 |
| 40 | Asselian | 298.9000 | 293.5200 | 5.3800 |
| 41 | Sakmarian | 293.5200 | 290.1000 | 3.4200 |
| 42 | Artinskian | 290.1000 | 283.5000 | 6.6000 |
| 43 | Kungurian | 283.5000 | 273.0100 | 10.4900 |
| 44 | Roadian | 273.0100 | 266.9000 | 6.1100 |
| 45 | Wordian | 266.9000 | 264.2800 | 2.6200 |
| 46 | Capitanian | 264.2800 | 259.5100 | 4.7700 |
| 47 | Wuchiapingian | 259.5100 | 254.1400 | 5.3700 |
| 48 | Changhsingian | 254.1400 | 251.9020 | 2.2380 |
| 49 | Induan | 251.9020 | 251.2000 | 0.7020 |
| 50 | Olenekian | 251.2000 | 247.2000 | 4.0000 |
| 51 | Anisian | 247.2000 | 242.0000 | 5.2000 |

| | | | | |
|---|---|---|---|---|
| 52 | Ladinian | 242.0000 | 237.0000 | 5.0000 |
| 53 | Carnian | 237.0000 | 227.0000 | 10.0000 |
| 54 | Norian | 227.0000 | 208.5000 | 18.5000 |
| 55 | Rhaetian | 208.5000 | 201.4000 | 7.1000 |
| 56 | Hettangian | 201.4000 | 199.5000 | 1.9000 |
| 57 | Sinemurian | 199.5000 | 192.9000 | 6.6000 |
| 58 | Pliensbachian | 192.9000 | 184.2000 | 8.7000 |
| 59 | Toarcian | 184.2000 | 174.7000 | 9.5000 |
| 60 | Aalenian | 174.7000 | 170.9000 | 3.8000 |
| 61 | Bajocian | 170.9000 | 168.2000 | 2.7000 |
| 62 | Bathonian | 168.2000 | 165.3000 | 2.9000 |
| 63 | Callovian | 165.3000 | 161.5000 | 3.8000 |
| 64 | Oxfordian | 161.5000 | 154.8000 | 6.7000 |
| 65 | Kimmeridgian | 154.8000 | 149.2000 | 5.6000 |
| 66 | Tithonian | 149.2000 | 145.0000 | 4.2000 |
| 67 | Berriasian | 145.0000 | 139.8000 | 5.2000 |
| 68 | Valanginian | 139.8000 | 132.6000 | 7.2000 |
| 69 | Hauterivian | 132.6000 | 125.7700 | 6.8300 |
| 70 | Barremian | 125.7700 | 121.4000 | 4.3700 |
| 71 | Aptian | 121.4000 | 113.0000 | 8.4000 |
| 72 | Albian | 113.0000 | 100.5000 | 12.5000 |
| 73 | Cenomanian | 100.5000 | 93.9000 | 6.6000 |
| 74 | Turonian | 93.9000 | 89.8000 | 4.1000 |
| 75 | Coniacian | 89.8000 | 86.3000 | 3.5000 |
| 76 | Santonian | 86.3000 | 83.6000 | 2.7000 |
| 77 | Campanian | 83.6000 | 72.1000 | 11.5000 |
| 78 | Maastrichtian | 72.1000 | 66.0000 | 6.1000 |
| 79 | Danian | 66.0000 | 61.6000 | 4.4000 |

| | | | | |
|---|---|---|---|---|
| 80 | Selandian | 61.6000 | 59.2000 | 2.4000 |
| 81 | Thanetian | 59.2000 | 56.0000 | 3.2000 |
| 82 | Ypresian | 56.0000 | 47.8000 | 8.2000 |
| 83 | Lutetian | 47.8000 | 41.2000 | 6.6000 |
| 84 | Bartonian | 41.2000 | 37.7100 | 3.4900 |
| 85 | Priabonian | 37.7100 | 33.9000 | 3.8100 |
| 86 | Rupelian | 33.9000 | 27.8200 | 6.0800 |
| 87 | Chattian | 27.8200 | 23.0300 | 4.7900 |
| 88 | Aquitanian | 23.0300 | 20.4400 | 2.5900 |
| 89 | Burdigalian | 20.4400 | 15.9800 | 4.4600 |
| 90 | Langhian | 15.9800 | 13.8200 | 2.1600 |
| 91 | Serravallian | 13.8200 | 11.6300 | 2.1900 |
| 92 | Tortonian | 11.6300 | 7.2460 | 4.3840 |
| 93 | Messinian | 7.2460 | 5.3330 | 1.9130 |
| 94 | Zanclean | 5.3330 | 3.6000 | 1.7330 |
| 95 | Piacenzian | 3.6000 | 2.5800 | 1.0200 |
| 96 | Pleistocene | 2.5800 | 0.0117 | 2.5683 |
| 97 | Holocene | 0.0117 | 0.0000 | 0.0117 |

**Table S2:** Equal-area latitudinal bins used in this study. Bins are generated assumming a regular spheroid Earth model with a mean radius of ~6,371 km.

| Bin | Minimum (º) | Midpoint (º) | Maximum (º) | Area ($m^2$) | Proportion of Area |
|---|---|---|---|---|---|
| 1 | 41.810 | 65.905 | 90.000 | 8.5e+13 | 0.167 |
| 2 | 19.471 | 30.641 | 41.810 | 8.5e+13 | 0.167 |
| 3 | 0.000 | 9.736 | 19.471 | 8.5e+13 | 0.167 |
| 4 | -19.471 | -9.736 | 0.000 | 8.5e+13 | 0.167 |
| 5 | -41.810 | -30.641 | -19.471 | 8.5e+13 | 0.167 |
| 6 | -90.000 | -65.905 | -41.810 | 8.5e+13 | 0.167 |

**Table S3:** Counts of the latitudinal bin with peak estimated genus richness (coverage-based rarefaction) for each hemisphere, within each stage, and for each Global Plate Model. Within the table, N denotes Northern Hemisphere latitudinal bins and S denotes Southern Hemisphere latitudinal bins. Note that for some stages, no estimates of genus richness could be generated for entire hemispheres, resulting in no peak estimated richness.

| Global Plate Model | High (N) | Middle (N) | Low (N) | Low (S) | Middle (S) | High (S) |
|---|---|---|---|---|---|---|
| GOLONKA | 11 | 46 | 38 | 37 | 38 | 21 |
| MERDITH2021 | 6 | 40 | 50 | 34 | 40 | 21 |
| PALEOMAP | 18 | 36 | 42 | 33 | 35 | 28 |
| TorsvikCocks2017 | 13 | 35 | 46 | 35 | 44 | 16 |

### 18 Supplementary figures

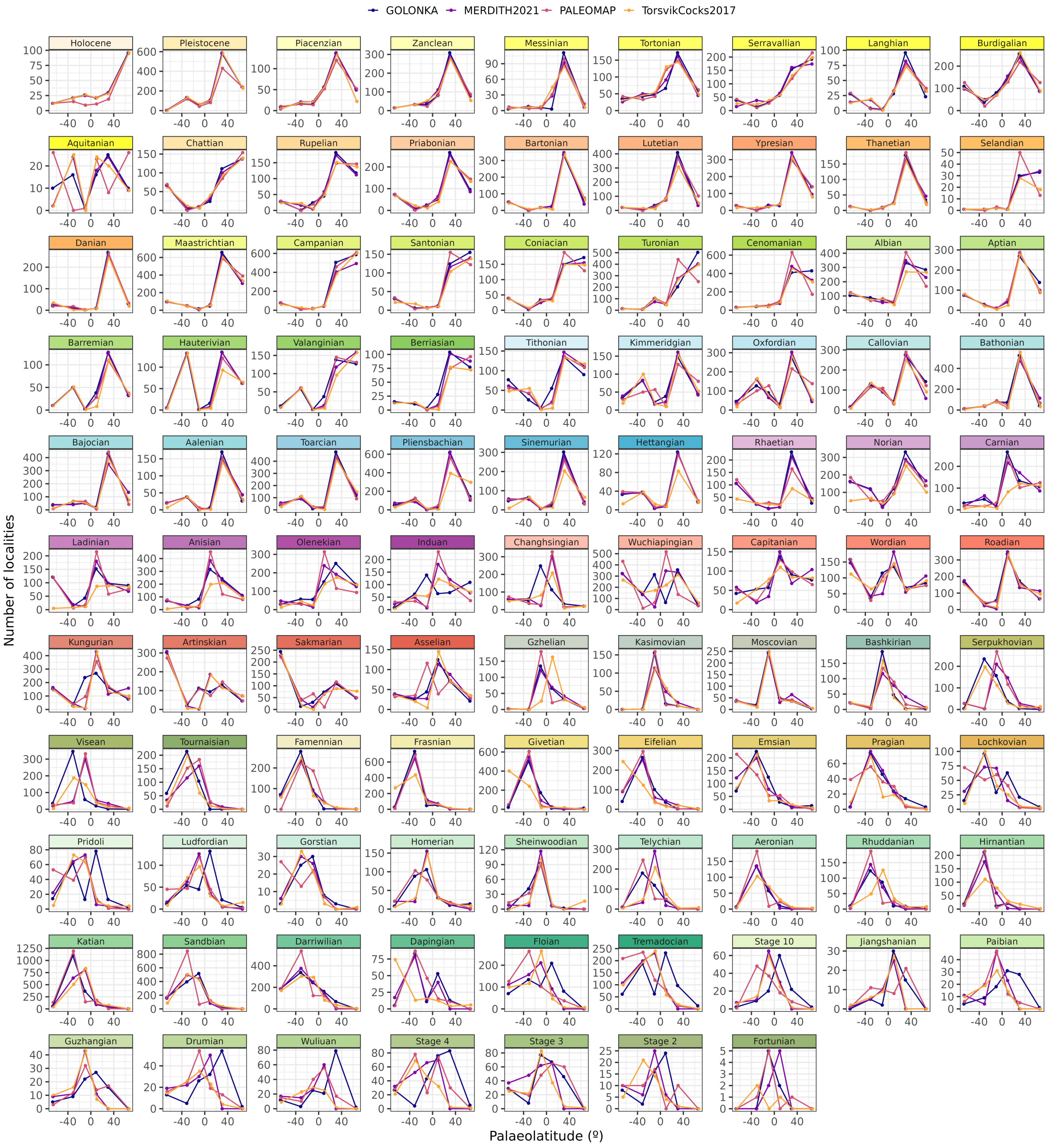


**Figure S1:** Number of fossil localities per palaeolatitudinal bin within each Phanerozoic stage that could be palaeogeographically reconstructed using four Global Plate Models: GOLONKA [1], PALEOMAP [2], MERDITH2021 [3], and TorsvikCocks2017 [4]. Differences in the number of localities within each palaeolatitudinal bin reflects differences between Global Plate Models in their palaeogeographic reconstruction.

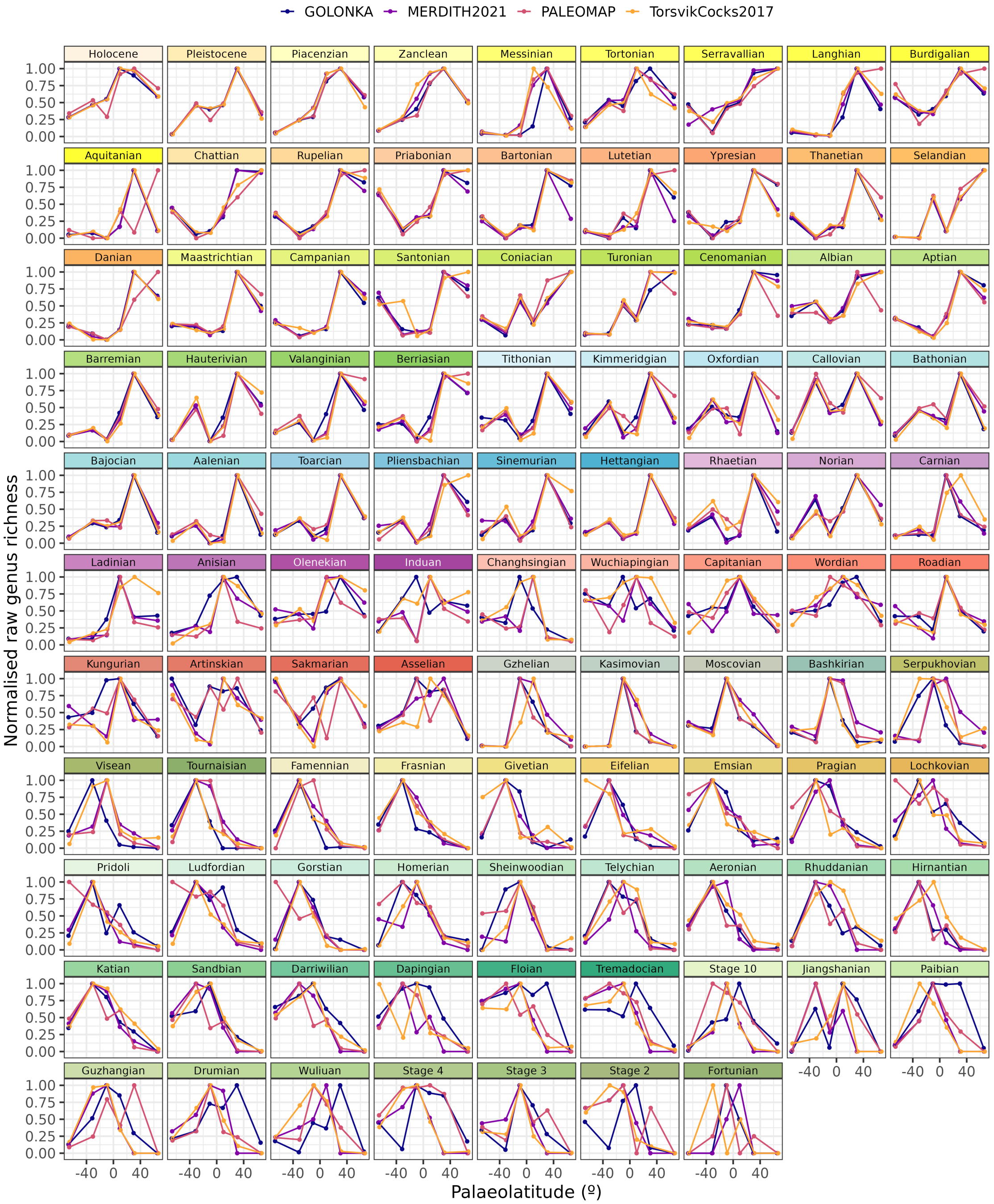


**Figure S2:** Phanerozoic stage-level reconstructions of the latitudinal biodiversity gradient for five major marine invertebrate groups (Bivalvia, Brachiopoda, Cephalopoda, Gastropoda, and Trilobita). Each individual plot depicts the raw normalised genus richness within each palaeolatitudinal bin for four Global Plate Models: GOLONKA [1], PALEOMAP [2], MERDITH2021 [3], and TorsvikCocks2017 [4]. Raw genus richness was normalised for each stage and Global Plate Model by dividing the genus counts within each palaeolatitudinal bin by the maximum value across palaeolatitudinal bins.

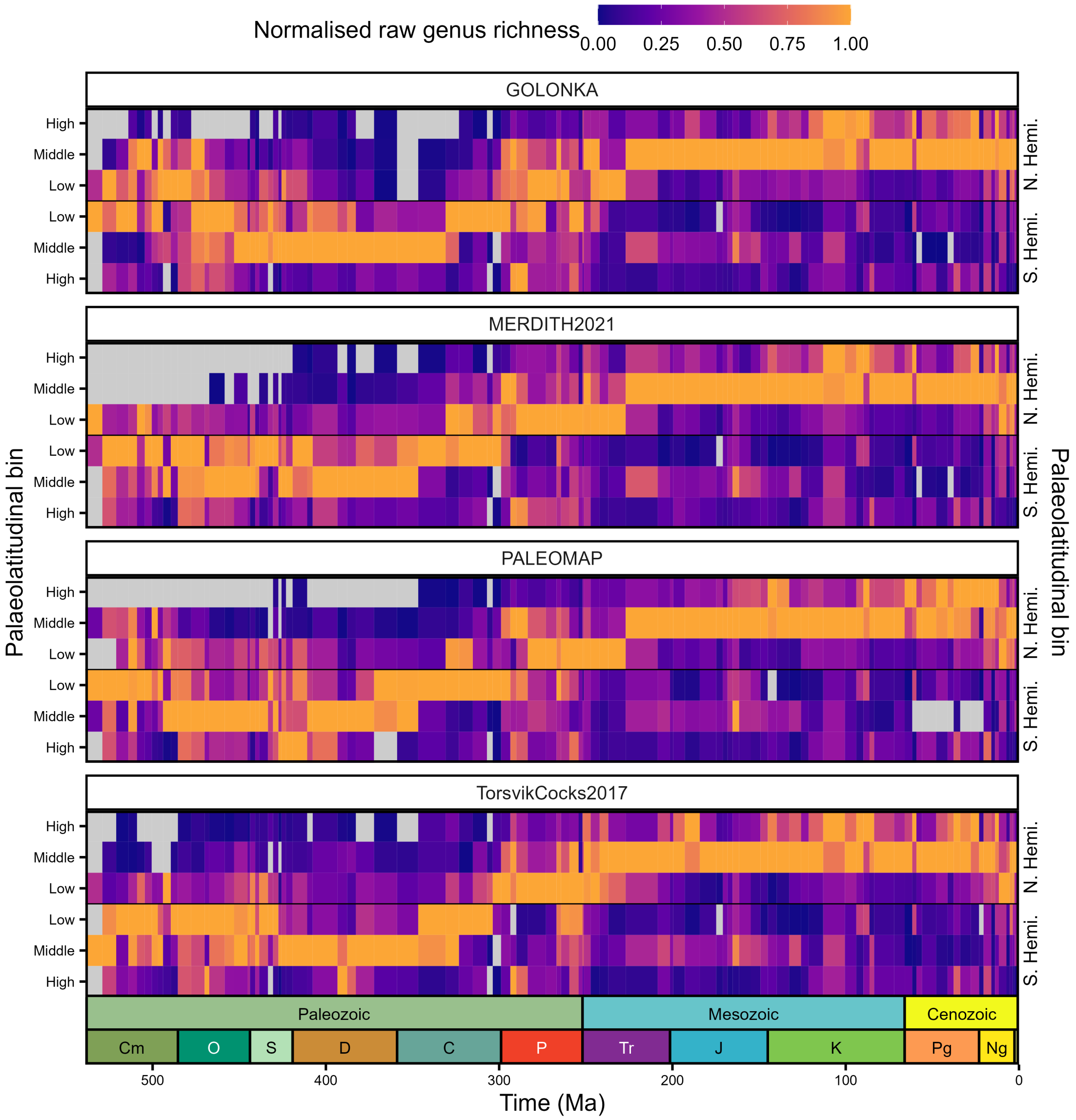


**Figure S3:** Phanerozoic heatmaps of raw genus counts for five major marine invertebrate groups (Bivalvia, Brachiopoda, Cephalopoda, Gastropoda, and Trilobita). Each individual panel depicts the normalised genus richness for four Global Plate Models: GOLONKA [1], PALEOMAP [2], MERDITH2021 [3], and TorsvikCocks2017 [4]. Heatmaps depict the normalised raw genus counts within palaeolatitudinal bins (low, middle, and high) within the Northern and Southern Hemisphere. Genus richness was normalised for each stage and Global Plate Model by dividing the genus richness within each palaeolatitudinal bin by the maximum value across palaeolatitudinal bins. Grey tiles indicate palaeolatitudinal bins without sufficient data to generate richness estimates. Period abbreviations are as follows: Cambrian (Cm); Ordovician (O), Silurian (S), Devonian (D), Carboniferous (C), Permian (P), Triassic (Tr), Jurassic (J), Cretaceous (K), Paleogene (Pg) and Neogene (Ng). The Quaternary is not depicted. The geological time scale axis was added to the plot using the R package 'deeptime' ver. 2.1.0 [5].

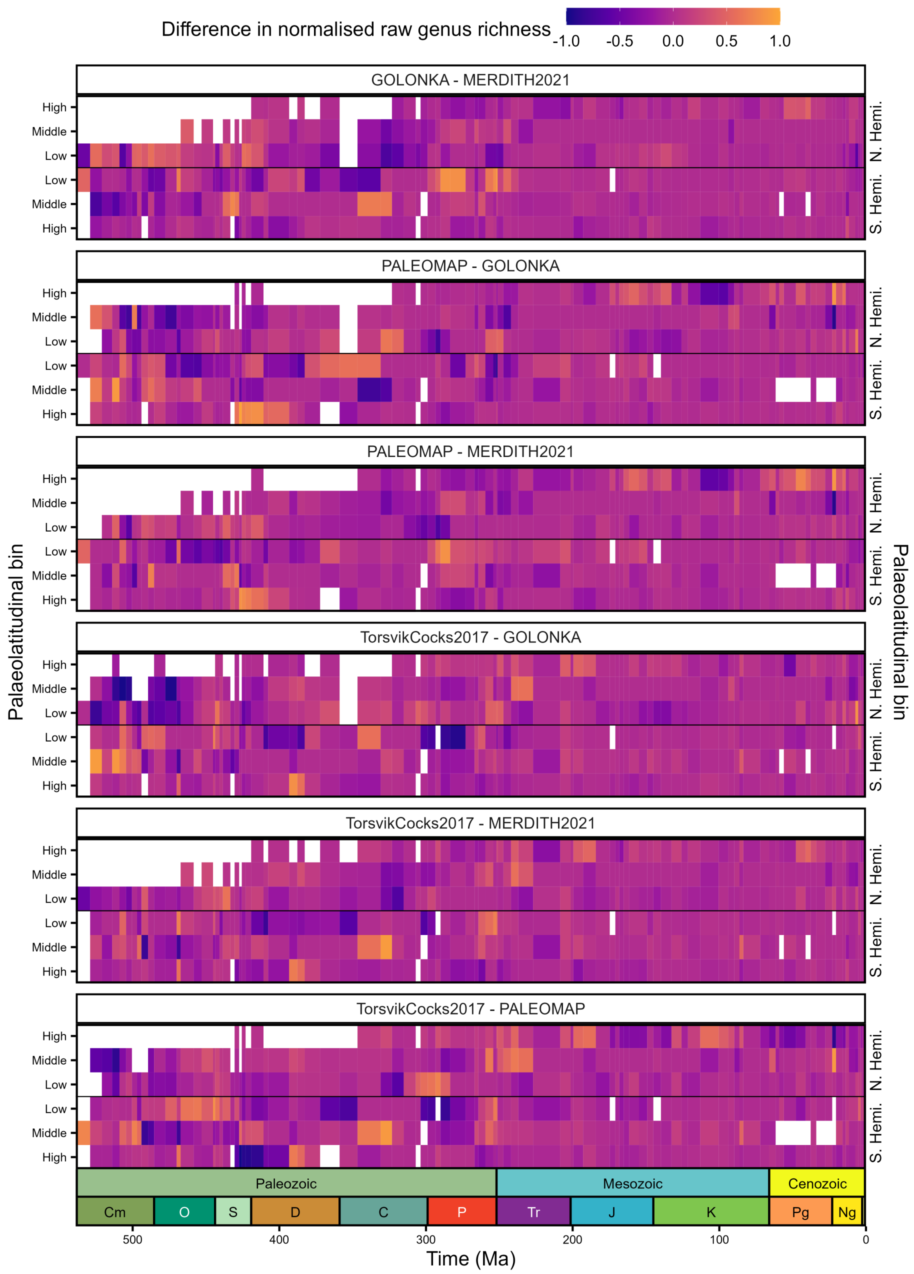


**Figure S4:** Phanerozoic heatmaps of the pairwise difference in normalised raw genus richness for four Global Plate Models: GOLONKA [1], PALEOMAP [2], MERDITH2021 [3], and TorsvikCocks2017 [4]. Genus richness was computed for each stage, palaeolatitudinal bin, and model using raw counts. Genus richness was normalised for each stage and Global Plate Model by dividing the genus richness within each palaeolatitudinal bin by the maximum value across palaeolatitudinal bins. White tiles indicate palaeolatitudinal bins without sufficient data to generate richness estimates or where estimates are not available for both being compared. Period abbreviations are as follows: Cambrian (Cm); Ordovician (O), Silurian (S), Devonian (D), Carboniferous (C), Permian (P), Triassic (Tr), Jurassic (J), Cretaceous (K), Paleogene (Pg) and Neogene (Ng). The Quaternary is not depicted. The geological time scale axis was added to the plot using the R package 'deeptime' ver. 2.1.0 [5].

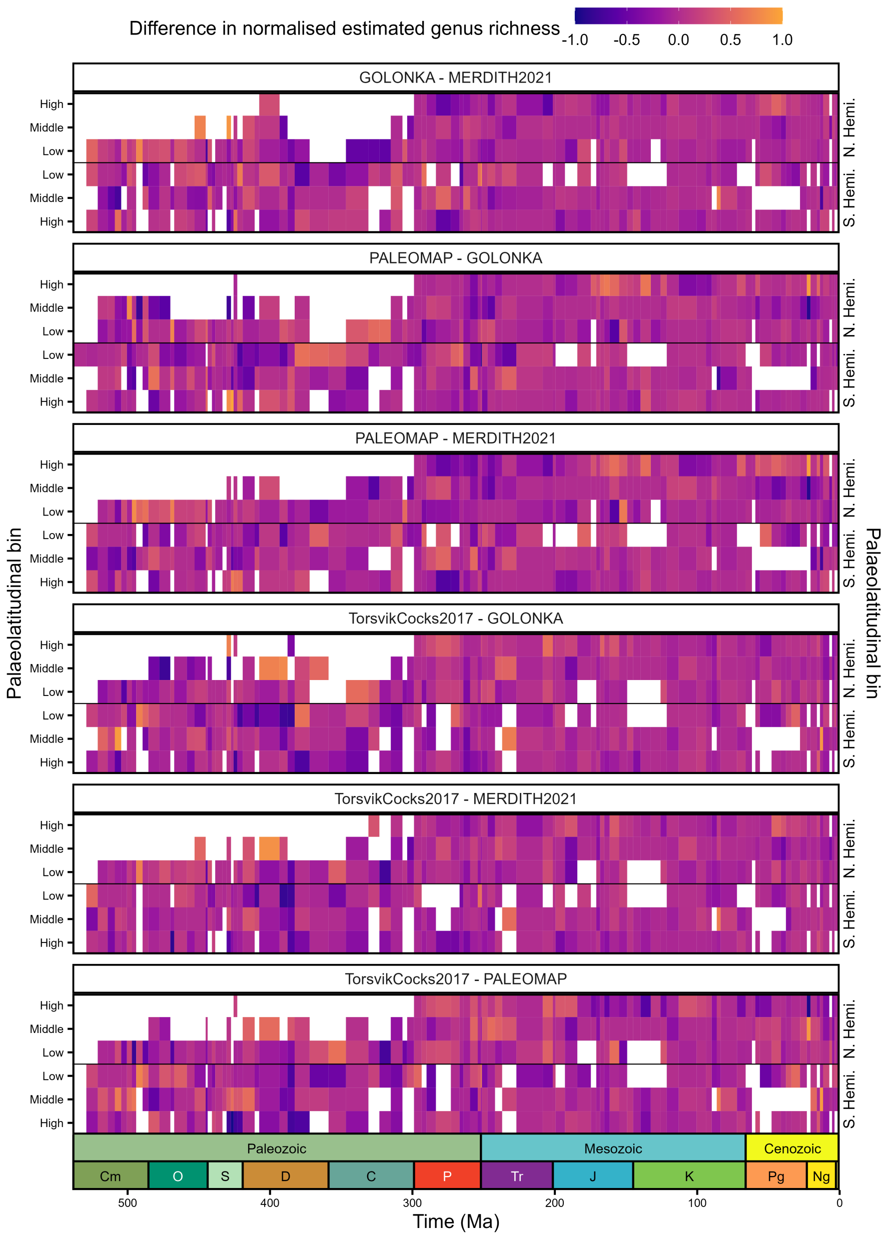


**Figure S5:** Phanerozoic heatmaps of the pairwise difference in normalised estimated (coverage-based rarefaction) genus richness for four Global Plate Models: GOLONKA [1], PALEOMAP [2], MERDITH2021 [3], and TorsvikCocks2017 [4]. Genus richness was estimated for each stage, palaeolatitudinal bin, and model using coverage-based rarefaction using the iNEXT R package ver. 3.0.1 [6] with a coverage level (otherwise known as a quorum level) of 0.4. Estimated genus richness was normalised for each stage and Global Plate Model by dividing the genus richness within each palaeolatitudinal bin by the maximum value across palaeolatitudinal bins. White tiles indicate palaeolatitudinal bins without sufficient data to generate richness estimates or where estimates are not available for both Global Plate Models being compared. Period abbreviations are as follows: Cambrian (Cm); Ordovician (O), Silurian (S), Devonian (D), Carboniferous (C), Permian (P), Triassic (Tr), Jurassic (J), Cretaceous (K), Paleogene (Pg) and Neogene (Ng). The Quaternary is not depicted. The geological time scale axis was added to the plot using the R package 'deeptime' ver. 2.1.0 [5].

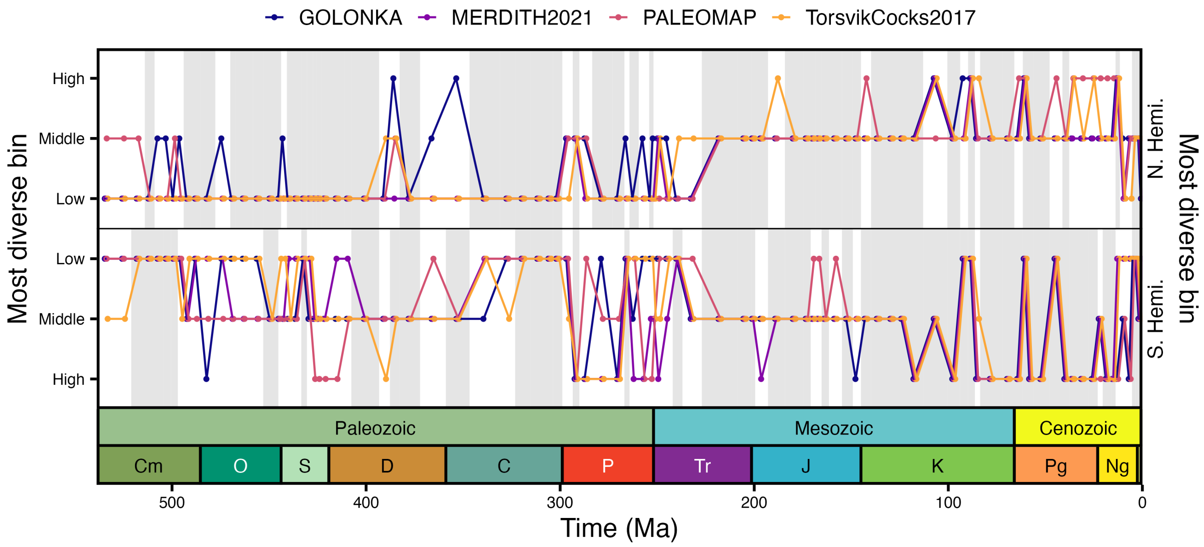


**Figure S6:** Phanerozoic plot of the palaeolatitudinal bin (low, middle, high) with the greatest raw genus counts within the Northern Hemisphere and Southern Hemisphere for each Global Plate Model: GOLONKA [1], PALEOMAP [2], MERDITH2021 [3], and TorsvikCocks2017 [4]. Grey shading highlights intervals in which the palaeolatitudinal bin with the greatest genus richness is in agreement across the four Global Plate Models. Period abbreviations are as follows: Cambrian (Cm); Ordovician (O), Silurian (S), Devonian (D), Carboniferous (C), Permian (P), Triassic (Tr), Jurassic (J), Cretaceous (K), Paleogene (Pg) and Neogene (Ng). The Quaternary is not depicted. The geological time scale axis was added to the plot using the R package 'deeptime' ver. 2.1.0 [5].

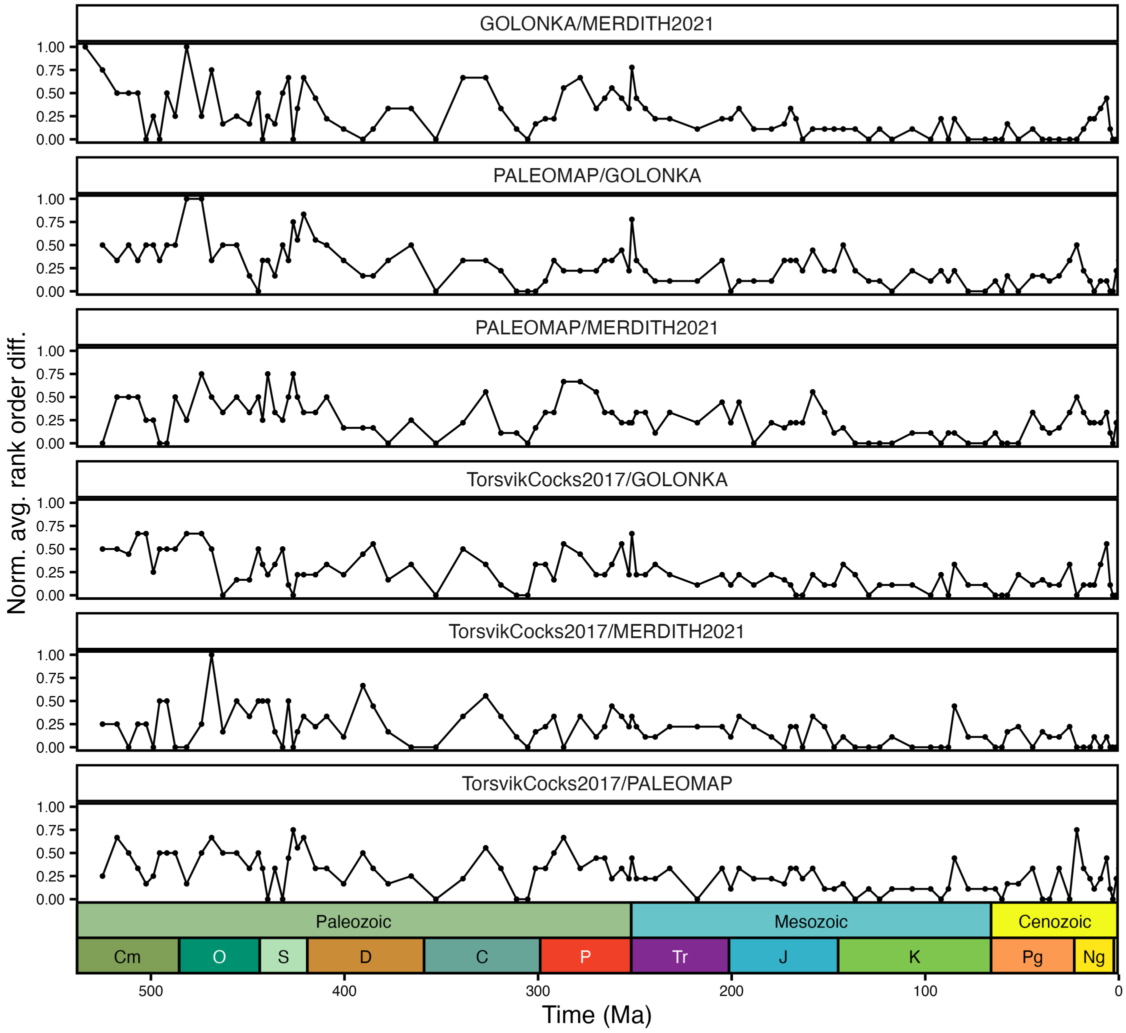


**Figure S7:** Phanerozoic plot of the normalised average rank order pairwise difference between latitudinal biodiversity gradients (raw genus richness) for each Global Plate Models: GOLONKA [1], PALEOMAP [2], MERDITH2021 [3], and TorsvikCocks2017 [4]. Average rank order differences are normalised by maximum possible average difference given the number of ranks available. Period abbreviations are as follows: Cambrian (Cm); Ordovician (O), Silurian (S), Devonian (D), Carboniferous (C), Permian (P), Triassic (Tr), Jurassic (J), Cretaceous (K), Paleogene (Pg) and Neogene (Ng). The Quaternary is not depicted. The geological time scale axis was added to the plot using the R package 'deeptime' ver. 2.1.0 [5].

19
